# Variational autoencoders learn universal latent representations of metabolomics data

**DOI:** 10.1101/2021.01.14.426721

**Authors:** Daniel P. Gomari, Annalise Schweickart, Leandro Cerchietti, Elisabeth Paietta, Hugo Fernandez, Hassen Al-Amin, Karsten Suhre, Jan Krumsiek

## Abstract

Dimensionality reduction approaches are commonly used for the deconvolution of high-dimensional metabolomics datasets into underlying core metabolic processes. However, current state-of-the-art methods are widely incapable of detecting nonlinearities in metabolomics data. Variational Autoencoders (VAEs) are a deep learning method designed to learn nonlinear latent representations which generalize to unseen data. Here, we trained a VAE on a large-scale metabolomics population cohort of human blood samples consisting of over 4,500 individuals. We analyzed the pathway composition of the latent space using a global feature importance score, which showed that latent dimensions represent distinct cellular processes. To demonstrate model generalizability, we generated latent representations of unseen metabolomics datasets on type 2 diabetes, schizophrenia, and acute myeloid leukemia and found significant correlations with clinical patient groups. Taken together, we demonstrate for the first time that the VAE is a powerful method that learns biologically meaningful, nonlinear, and universal latent representations of metabolomics data.

## 1. Introduction

Modern metabolomics experiments yield high-dimensional datasets with hundreds to thousands of measured metabolites in large human studies with thousands of participants^1^. Such datasets are routinely generated to profile the molecular phenotype of disease and identify the underlying pathological mechanisms of action^2–5^. Extracting systemic effects from high-dimensional datasets requires dimensionality reduction approaches to untangle the high number of metabolites into the processes in which they participate. To this end, linear dimensionality reduction methods, such as principal component analysis (PCA) and independent component analysis (ICA), have been extensively applied to high-dimensional biological data^6–10^. However, metabolic systems, like most complex biological processes, contain nonlinear effects which arise due to high-order enzyme kinetics and upstream gene regulatory processes^11,12^. For example, metabolite ratios are an intuitive and widely used approach to detect nonlinear effects in metabolomics data, approximating the steady state between reactants and products of metabolic reactions^13,14^. Extending this concept, systematic methods that take nonlinearities into account are required to correctly recover the functional interplay between metabolites in an unbiased fashion.

Autoencoders (AEs) are a type of neural network architecture developed as an unsupervised dimensionality reduction method that can capture nonlinear effects^15^. AEs reduce high-dimensional data into latent variables through an encoding/decoding process which recreates the input data after passing through a lower dimensional space. Once the model is fitted, the latent variables represent a compact, often easier-to-interpret version of the original data. While AEs have been successful for prediction tasks on biological datasets^16,17^, they tend to learn latent spaces specifically fitted to the input dataset and are therefore not generalizable to unseen data. To address this, Variational Autoencoders (VAEs) were introduced as a probabilistic extension of the AE architecture that constrains the latent variables to follow a predefined distribution^18^. With this extension, the VAE not only reconstructs the input data, but infers the generative process behind the data, leading to high generalizability across datasets. The VAE architecture has, for example, proven effective for predicting cell-level response to infection in transcriptomic data not available during training, and predicting drug response from gene expression data where drug response information is sparse^19–22^.

The application of deep learning architectures to metabolomics datasets has significantly lagged behind all other omics^23^ due to the unavailability of large metabolomics cohorts. By applying VAE architectures to metabolomics data, we have the potential to learn more accurate latent dimension representations that take nonlinearities into account. In addition, the probabilistic structure of VAEs will learn latent dimensions that are generalizable across multiple datasets.

In this paper, we trained a VAE model on 217 metabolite measurements in 4,644 blood samples from the TwinsUK study^24^ and evaluated our model performance in comparison to a linear PCA model (Figure 1a). To investigate the biological relevance of the learned VAE and PCA latent dimensions, we employed the Shapley Additive Global Importance (SAGE) method^25^, which determines the contribution of each input to each latent dimension. We calculated SAGE values at different granularities, i.e., metabolites, *sub-pathways*, and *super-pathways* (Figure 1b). We then applied the models on three additional blood metabolomics datasets to test their ability to recover disease phenotypes in unseen datasets: Type 2 Diabetes diagnosis in The Qatar Metabolomics Study on Diabetes (QMDiab, *n* = 358), therapy response in an acute myeloid leukemia dataset (AML, *n* = 85), and schizophrenia diagnosis in a third validation dataset (*n* = 207) (Figure 1c).

**Figure 1.**
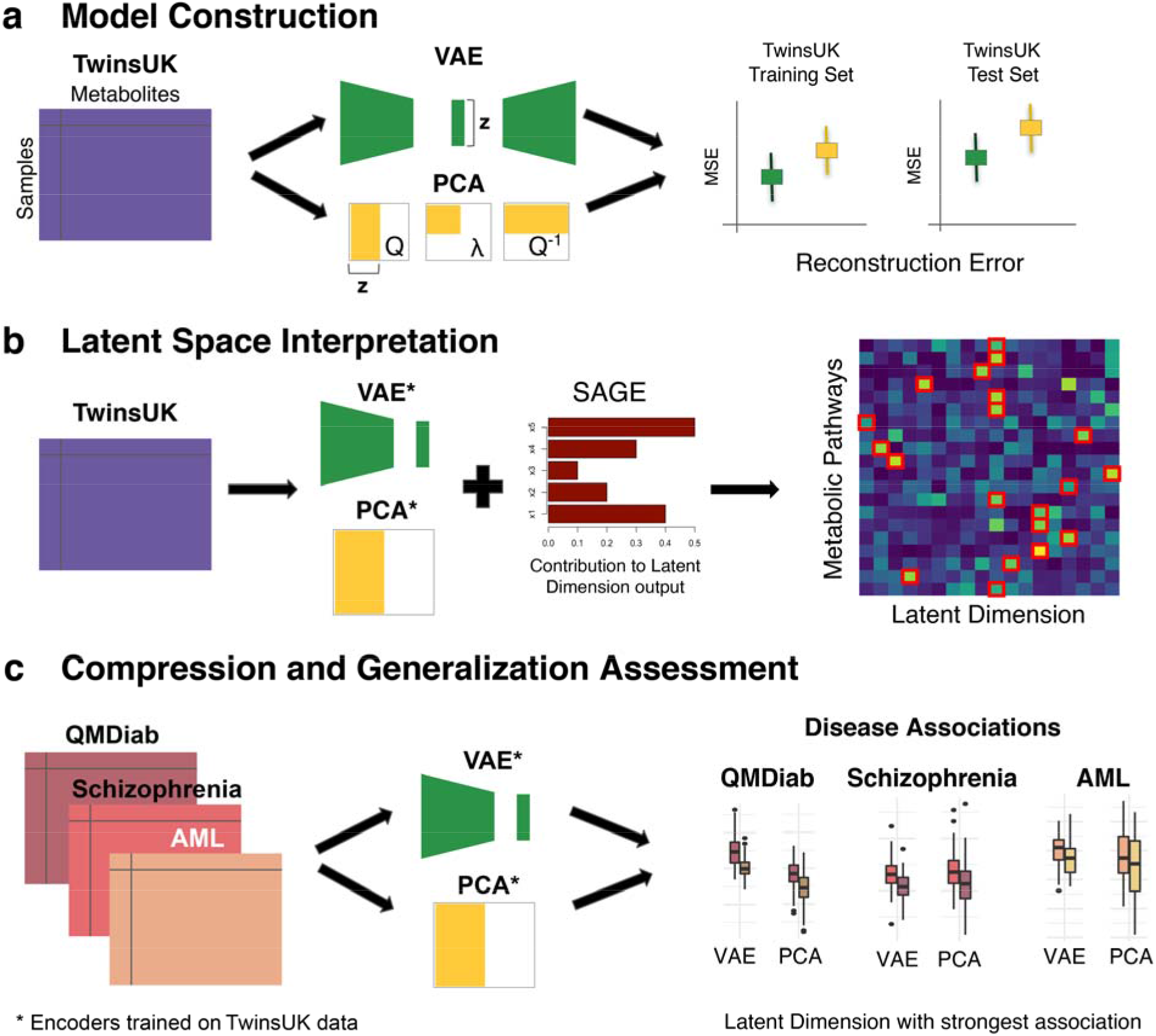
Overview of our approach. **a**, VAE and PCA models were trained using training and test sets in the TwinsUK dataset (*n*=4,644 samples, *p*=217 metabolites). Model performance was then evaluated using Mean Squared Error (MSE) of metabolite correlation matrix reconstruction. **b**, The SAGE method was applied to calculate the contribution of individual metabolites, sub-pathways and super-pathways to each latent dimension. **c**, QMDiab (*n* = 358), AML (*n* = 85), and Schizophrenia (*n* = 207) datasets were encoded using VAE and PCA models trained on the TwinsUK data. Latent dimensions of each model were then associated with disease phenotypes.

## 2. Results

### 2.1. VAE model construction and fitting

Our VAE architecture consisted of an input/output layer, an intermediate layer and a latent layer. We split the TwinsUK cohort into an 85% training and a 15% test set, and the training set was used to optimize the hyperparameters in the VAE model. Keras Tuner^26^ identified the following optimal hyperparameters: Intermediate layer dimensionality = 200, learning rate = 0.001, and Kullback-Leibler (KL) divergence weight = 0.01. With these parameters fixed, we optimized the dimensionality *d* of the latent layer **z** by calculating the reconstruction MSE of the correlation matrix (CM-MSE) of metabolites (Figures 2a and 2b). We observed that the CM-MSE curve plateaus after *d* = 18, indicating that increasing the latent dimensionality beyond this value only marginally improves the models. The final architecture of the model consisted of a 217-dimensional input/output layer (the number of metabolites in our datasets), a 200-dimensional intermediate layer, and an 18-dimensional latent layer (Figure 2c).

**Figure 2.**
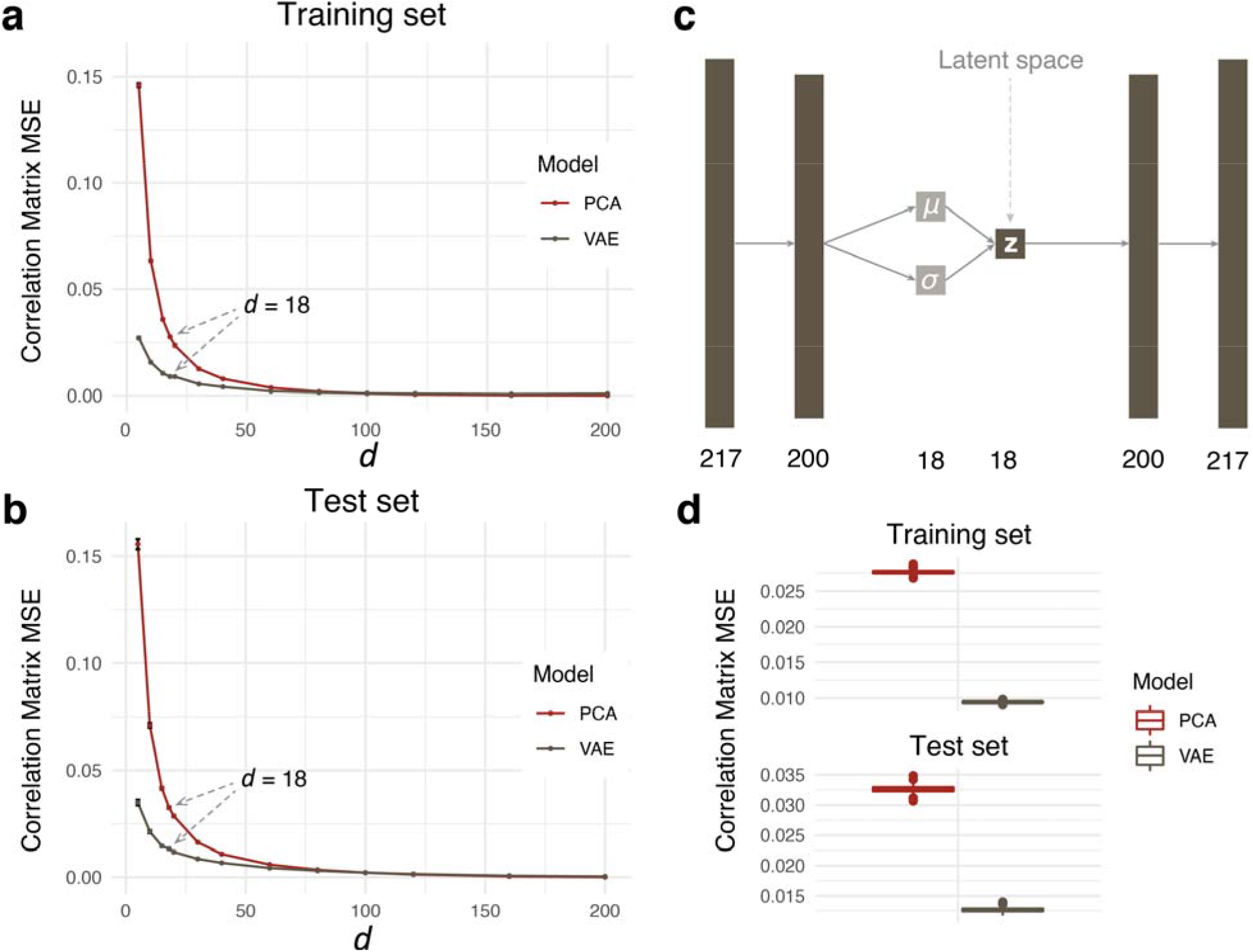
VAE and PCA model construction on the TwinsUK dataset. **a**, Training and **b**, test set metabolite correlation matrix reconstruction for a range of latent dimensionality values *d*. The slope of the VAE curve plateaus after *d* = 18. Error bars correspond to one standard deviation from bootstrapping. **c**, Final VAE architecture, where ◻ is the mean vector and ◻ is the standard deviation vector that generates the latent space **z**. **d**, Reconstruction MSE for latent dimensionality *d* = 18 on training (top) and test sets (bottom). The VAE preserved feature correlations substantially better than PCA.

We used principal component analysis (PCA) as a baseline model to compare the VAE to a linear latent variable embedding method. To this end, we fitted a PCA on the TwinsUK train data and extracted the first *d* = 18 dimensions, i.e., principal components. While PCA reconstructs the data matrix better than the VAE (Extended Data Figure 1), the VAE outperforms PCA in terms of correlation matrix reconstruction via CM-MSE in both the TwinsUK train and test set (Figure 2d).

This discrepancy between the MSE on the correlation matrix and the more commonly used sample-wise MSE^18^, where PCA outperforms our model, outlines ambiguities in the methods to assess VAE reconstruction performances. Notably, other authors have shown previously that sample reconstruction performance does not necessarily imply better model performance^20,22^. Our results suggest that while the VAE does not reconstruct the original data matrix precisely, it is superior to PCA at preserving metabolite correlations.

### 2.2. Interpretation of VAE latent space dimensions in the context of metabolites and pathways

We evaluated the composition of all latent dimensions in the context of metabolic pathways. For each metabolite in our dataset, a “sub-pathway” and “super-pathway” annotation was available (see Methods 4.2.). Sub-pathways refer to biochemical processes such as “TCA Cycle” and “Sphingolipid Metabolism”, while super-pathways are broad groups such as “Lipid” and “Amino acid”. To provide insights into the processes represented by different VAE dimensions, we computed SAGE scores, a measure of model feature relevance, at the level of metabolites, sub-pathways and super-pathways (Figure 3 and Extended Data Figure 2 for VAE, Extended Data Figure 3 for PCA).

**Figure 3.**
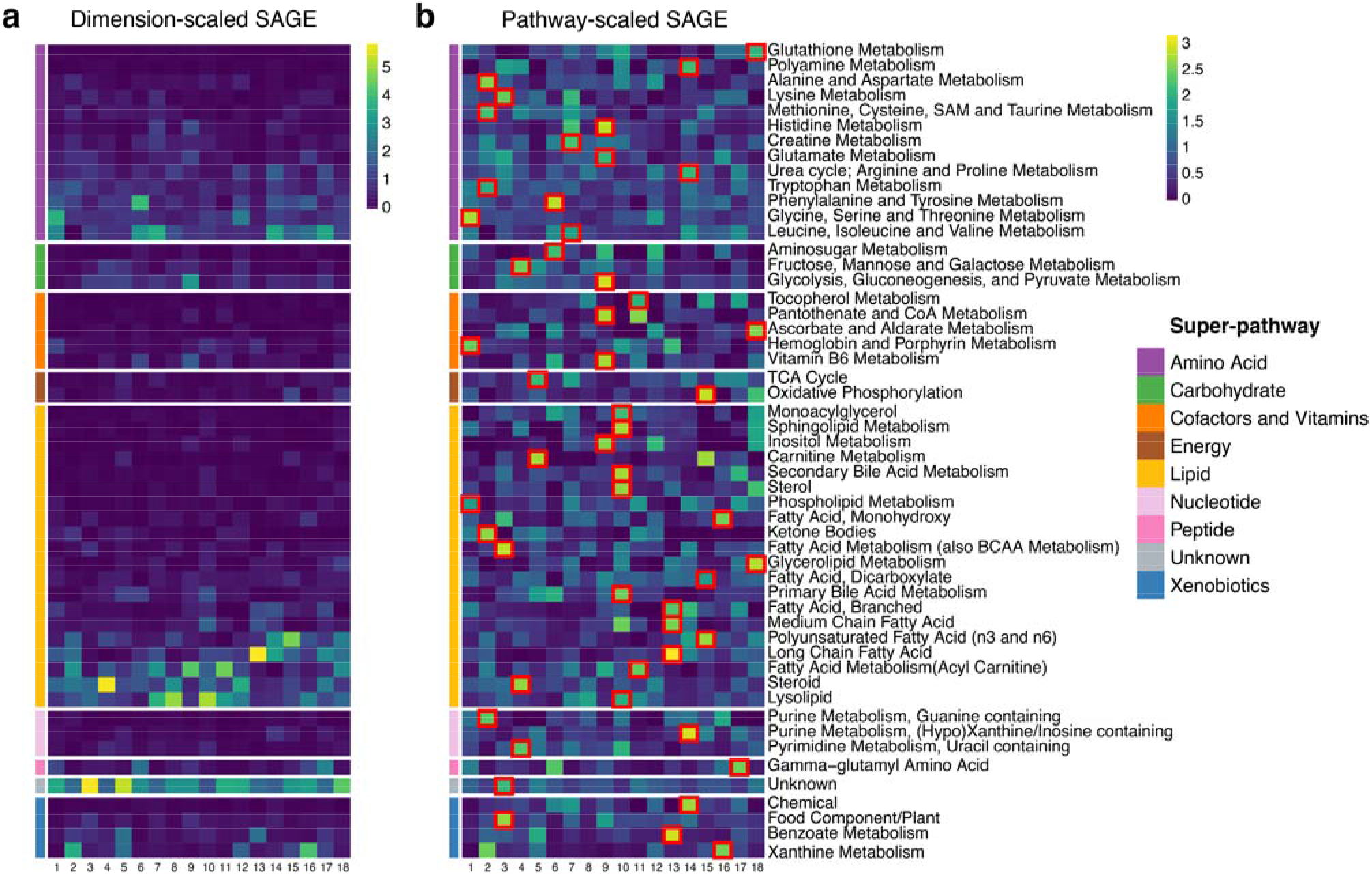
Sub-pathway-level SAGE values for the VAE latent dimensions. **a**, SAGE values were scaled by dimension, i.e., set to standard deviation 1 for each column in the matrix. This highlights pathways that contribute the most to each dimension. Lipid and amino acid super-pathways showed the highest values for most dimensions, which can be attributed to the high number of metabolites in those pathways. **b**, SAGE values were scaled by pathway, i.e., set to standard deviation 1 for each row in the matrix. This highlights dimensions that contribute to a pathway the most. Taking into consideration the largest scaled SAGE values per pathway (red square marks), almost all sub-pathways are represented by unique dimensions. The combination of these key sub-pathways of a dimension outlines the distinct cellular mechanisms a dimension encodes.

The VAE sub-pathway heatmap (Figure 3a) shows that nearly all dimensions have major contributions by lipid and amino acid super-pathways. The prevalence of the two super-pathways can be attributed to the fact that those groups contain the largest number of metabolites in the dataset. Note that we deliberately omitted the “Unknown” molecule group, which refers to unidentified metabolites that could originate from any pathway.

Inspecting the SAGE values in the other direction, almost all sub-pathways are predominantly represented by a single VAE dimension that captures the respective pathway the most (Figure 3b, red square marks). For instance, “glycolysis, gluconeogenesis and pyruvate metabolism” and other functionally related sub-pathways of central carbon metabolism are represented by VAE dimension 9. Another interesting example is VAE dimension 15, which captures functionally-related essential mitochondrial processes, such as oxidative phosphorylation, dicarboxylic fatty acids, and n3 and n6 polyunsaturated fatty acid metabolism. Taken together, these results show that VAE latent dimensions capture a complex mix of functionally-related sub-pathways, thus capturing major metabolic processes in the dataset.

In contrast, PCA dimensions 1 to 3, which by construction represent the highest linear variations in the data, nonspecifically capture various sub-pathways. Most other PCA dimensions also primarily contain unrelated sub-pathways (Extended Data Figure 3).

### 2.3. VAE latent space captures signals in unseen diabetes, schizophrenia, and cancer metabolomics datasets

We investigated whether VAE latent dimensions learned on the TwinsUK data contained information that is generalizable to other datasets. To this end, we encoded metabolomics data from three clinical datasets, type 2 diabetes, schizophrenia, and acute myeloid leukemia (AML) using the VAE and PCA encoders trained on TwinsUK dataset. For each VAE and PCA latent dimension, we performed a two-sided t-test between diabetic vs. non-diabetic individuals, schizophrenic vs. non-schizophrenic individuals, and full vs. no response in an AML clinical trial, respectively. Across all datasets, the best performing VAE dimensions associated substantially stronger with the patient groups than any of the PCA dimensions (Figures 4a-f). The strength of associations between VAE dimensions and disease parameters were comparable to single metabolite associations (Extended Data Tables 1-3). However, unlike the VAE dimensions, these univariate associations do not represent system-level mechanisms related to the diseases.

**Figure 4.**
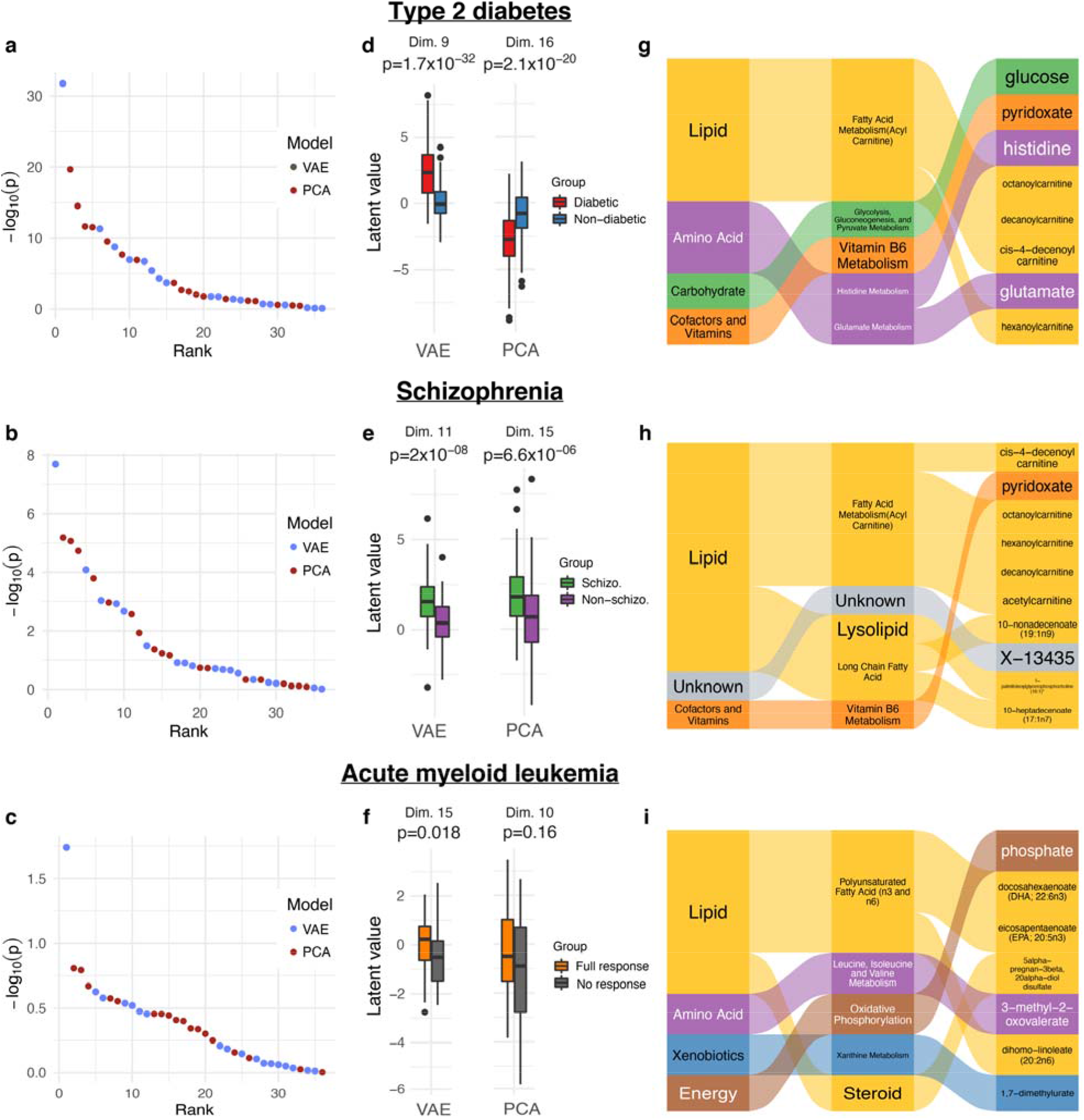
VAE latent space associations with clinical outcomes. **a**, **b**, **c,** Sorted -log_10_(p-value) for all VAE and PCA dimensions for the type 2 diabetes, schizophrenia and AML datasets, respectively. The highest scoring VAE dimensions showed considerably lower p-values than the highest scoring PCA dimensions for all datasets. **d**, **e**, **f,** Latent space dimensions with the lowest p-values for the three datasets. **g**, **h**, **i,** Contributions of super-pathways, sub-pathways and metabolites to the highest scoring VAE latent dimensions, determined by SAGE values. All dimensions are driven by lipid metabolism and a mixture of other super-pathways, with differing sub-pathways contributing to the different dimensions. p = p-value. Schizo. = schizophrenic.

To obtain a better understanding of the driving factors of the VAE associations, we ranked pathways and metabolites by their calculated SAGE values (Figures 4g-i):

#### Type 2 diabetes

VAE latent dimension 9 showed the highest association with type 2 diabetes, with a considerably stronger signal than the highest correlating PCA dimension 16 (p=1.7×10^−32^ vs. p=2.1×10^−20^, respectively; Figure 4d). The top sub-pathways were “acyl carnitine fatty acid metabolism”, “glycolysis, gluconeogenesis, and pyruvate metabolism”, “vitamin B6 metabolism”, and “histidine metabolism”. The top-ranking metabolite in dimension 9 was glucose, which is directly affected by the disease and thus serves as a positive control. Other high-ranking metabolites included pyridoxate, histidine, and medium chain acyl-carnitines (Figure 4g). Vitamin B6 metabolism, which includes pyridoxate, has been shown to associate with type 2 diabetes and with the predisposition of diabetic patients to other diseases^27,28^. Additionally, circulating medium chain acyl-carnitines have been shown to be associated with early stages of type 2 diabetes^29,30^. We furthermore correlated dimension 9 with clinical lab measurements from the QMDiab study and found a strong association between this dimension and HbA1c (p=5.6×10^−56^ compared to PCA p=1.1×10^−30^, Extended Data Figure 4), a widely used diabetes biomarker^31,32^. This finding demonstrates how a quantitative disease biomarker can carry more information than a crude disease yes/no classification, and further highlights the higher information content in the VAE latent dimensions compared to PCA.

#### Schizophrenia

VAE dimension 11 had a stronger association with schizophrenia than PCA dimension 15 (p=2.0×10^−8^ vs. p=6.6×10^−6^, respectively; Figure 4e). The top scoring metabolites for this dimension (Figure 4h) were mainly acyl-carnitines, such as 4-decanoylcarninite, octanoylcarnitine, and hexanoylcarnitine, and a series of lysolipids. Acyl-carnitines, which are involved in energy metabolism and reflect an individual’s mitochondrial beta-oxidation function, have been previously shown to be associated with schizophrenia^33,34^. Vitamin B6 metabolism, through pyridoxate, is also one of the highest-ranking pathways for this dimension. Previous studies have demonstrated that low levels of vitamin B6 are associated with a subgroup of schizophrenic patients^35,36^.

#### Acute myeloid leukemia (AML)

AML response groups associated an order of magnitude stronger with VAE dimension 15 than with PCA dimension 10 (p=0.018 vs. p=0.16, respectively; Figure 4f). Note that the p-value would not withstand multiple testing correction (Extended Data Tables 1 and 2); the detected signal is thus merely suggestive and requires replication in future studies. Phosphate, which regulates the oxidative phosphorylation pathway and is involved in energy metabolism, is the most important metabolite for dimension 15. It has been previously demonstrated that oxidative phosphorylation plays a paramount role in AML survival and drug resistance^37–39^ and could be an effective target for combination therapy in chemoresistant AML^38,40^. Additionally, dimension 15 is driven by various metabolites from the n3 and n6 polyunsaturated fatty acid (PUFA) sub-pathway, such as docosahexaenoate (DHA) and eicosapentaenoate (EPA) (Figure 4i). It has been shown that treatment of AML cell lines with DHA and EPA has deleterious effects on their mitochondrial metabolism which leads to cell death^41–44^, indicating that PUFAs might play an essential role in AML. We furthermore investigated correlations of the latent dimensions with 21 major AML-related mutations; the analysis revealed no noteworthy results (Extended Data Figure 5).

Taken together, these results suggest that our VAE has learned representations of metabolic processes that are essential for unseen clinical outcomes.

## 3. Discussion

In this study, we trained a VAE on metabolomics data from the TwinsUK population cohort and applied the learned latent representations on unseen data. Our VAE model outperformed PCA in metabolite correlation matrix reconstruction. Interpretation of VAE latent dimensions at the metabolite, sub-pathway, and super-pathway level revealed that these dimensions represent functionally-related and distinct cellular processes. Moreover, VAE latent dimensions showed substantially stronger disease associations than PCA in unseen Type 2 Diabetes, schizophrenia, and AML datasets. This implies that the VAE learned a latent representation of metabolomics data that is biologically informative and transferable across different cohorts.

The generalizability of the VAE across different datasets is especially remarkable given the vastly different underlying populations of the datasets analyzed here. The VAE was trained on the TwinsUK population cohort, a European-ancestry population cohort consisting predominantly of British women (~92%), while the validation datasets are mixed-gender and multi-ethnic cohorts from the US and Qatar. Despite the existence of these variations in our datasets, our VAE learned a generalized representation of metabolomics data which was able to identify disease-related differences.

The main limitation of our study is the size of the TwinsUK training dataset with *n=*4,644. This is a general issue with human subject metabolomics studies, where even the largest cohorts reach only about *n=*15,000^45^. Deep learning models are currently more popular in larger datasets of *n=*60,000 samples or more, such as single cell transcriptomic^16,46–48^, image^49–51^, and text sources^52^. Learning the variation in such large datasets allows these models to significantly outperform their linear counterparts. Large metabolomics datasets, such as that of the UK BioBank with a sample size of up to *n =* 500,000^53^, will be available in the near future, and will enable the creation of more expressive and deeper VAE models.

To the best of our knowledge, this is the first study to construct a universal latent representation of metabolomics data using VAEs. Our results show that VAEs are well-suited for metabolomics data analysis and can potentially replace dimensionality reduction approaches, such as PCA, in creating a universal, systems-level understanding of metabolism.

## 4. Methods

### 4.1. Datasets

The TwinsUK registry is a population-based study of around 12,000 volunteer twins from all over the United Kingdom. The participants have been recruited since 1992 and are predominantly female, ranging in age from 18 to 103 years old. Study design, sampling methods, and data collection have been described elsewhere^24^. For our study, we included data from 4,644 twins (4,256 females, 388 males), the subset of TwinsUK for which plasma metabolomics measurements were available. Ethical approval was granted by the St Thomas’ Hospital ethics committee and all participants provided informed written consent.

The QMDiab study was conducted between February and June of 2012 at the Dermatology Department of Hamad Medical Corporation (HMC) in Doha, Qatar. The study population was between the ages of 23 and 71, predominantly of Arab, South Asian, and Filipino descent. Data collection and sampling methods have been previously described elsewhere^54^. For this study, we included plasma data of 358 subjects (176 females, 182 males; 188 diabetic, 177 non-diabetic). The study was approved by the Institutional Review Boards of HMC and Weill Cornell Medicine-Qatar (WCM-Q). Written informed consent was obtained from all participants.

For the schizophrenia analysis, metabolomics samples were taken from an antipsychotics study conducted in Qatar between December 2012 and June 2014^55^. A total of 226 participants between the ages of 18 and 65 years of age were recruited, predominantly of Arab descent. For our study, we included plasma metabolomics measurements from 207 subjects (84 females, 142 males; 102 schizophrenic, 105 non-schizophrenic). Approval for the study was obtained from the HMC and WCM-Q Institutional Review Boards, and all participants provided written informed consent.

The cohort of patients with acute myeloid leukemia (AML) comes from the ECOG-ACRIN Cancer Research Group phase 3 trial NCT00049517. This study was conducted between December 2002 and November 2008, recruiting 657 patients with AML between the ages of 17 and 60. A subset of these patients had follow-up profiling to determine their response to therapy. For this study, we included the serum metabolomics measurements of 85 subjects of which 43 responded to therapy and 42 did not (34 females, 51 males). The study was approved by the institutional review board at the National Cancer Institute and each of the study centers, and written informed consent was provided by all patients.

### 4.2. Metabolomics measurements and metabolite annotations

Metabolic profiling for all four cohorts was performed using non-targeted ultrahigh-performance liquid chromatography and gas chromatography separation, coupled with mass spectrometry on the Metabolon Inc. platform as previously described^56^. Notably, the AML dataset was based on serum samples, while TwinsUK, QMDiab, and schizophrenia metabolomics were run on plasma samples. However, previous studies have shown that these two sample types are comparable, as shown by high correlations and good reproducibility between plasma and serum measurements in the same blood sample^57^.

For each metabolite measured on the Metabolon platform, a super-pathway and sub-pathway annotation was provided. For super-pathways, we have nine annotations referring to broad biochemical classes, namely “Amino acid”, “Carbohydrate”, “Cofactors and vitamins”, “Energy”, “Lipid”, “Nucleotide”, “Peptide”, “Xenobiotics”, and “Unknown”. Note that “Unknown” is assigned to unidentified metabolites. Furthermore, we have 54 sub-pathway which represent more functional metabolic processes, such as “Carnitine metabolism”, “TCA Cycle”, and “Phenylalanine and Tyrosine Metabolism”.

### 4.3. Data processing and normalization across datasets

For each dataset, metabolite levels were scaled by their cohort medians, quotient normalized^58^ and then log-transformed. Samples with more than 30% missing metabolites and metabolites with more than 10% missing samples were removed. Missing values were imputed using a k-nearest neighbors imputation method^59^. Datasets with BMI measurements (Schizophrenia, QMDiab, and Twins) were corrected for that confounder and then mean-scaled. 217 metabolites were overlapping between the 4 datasets and were kept for further analysis.

Semi-quantitative, non-targeted metabolomics measurements are inherently challenging to compare across datasets due to heterogeneity between studies. This prevents any machine learning model from being transferable from one study to the other. To ensure comparability, datasets were normalized using a uniform group of participants as a reference set. This group was selected as follows: Male, within a 20-year age range (30-50 for TwinsUK, QMdiab, and schizophrenia, 40-60 for AML due to low sample size of younger participants), BMI between 25 and 30 (not available for AML data, thus not filtered for that dataset), and in the respective control group. Each metabolite in each dataset was then scaled by the mean and standard deviation of their respective uniform sample groups. The assumption of this approach is that the uniform group of reference participants has the same distributions of metabolite concentrations.

### 4.4. Variational Autoencoders

To train our VAE model, we first split the TwinsUK data into 85% training and 15% test sets. We then fixed our VAE architecture to be composed of an input/output layer, an intermediate layer which contains nonlinear activation functions, and a *d*-dimensional latent layer. The latent layer consists of a mean vector ◻ and a standard deviation vector ◻, both of length *d*, which parametrize each latent dimension as a Gaussian probability distribution. This latent space, denoted by ***z***, is constructed by the simultaneous learning of the ◻ and ◻ encoder through the use of a reparameterization trick that enables back propagation during training^18^. The *d* x *d* covariance matrix L of the underlying multivariate Gaussian is assumed to be diagonal (i.e., no correlation across latent dimensions), allowing the covariance matrix to be represented by a single vector ◻ of length d.

For the parameter fitting procedures, all weights were initialized using Keras’ default model weight initialization, i.e. Glorot uniform^60^. Leaky rectified linear units (ReLUs)^61^ were used for nonlinear activation functions. The VAE models were trained for 1,000 epochs using MSE loss for sample reconstruction and a batch size of 32.

To select the latent dimensionality *d* of our VAE model, we initially fixed this value to *d* = 50. We then optimized the model hyperparameters using Keras Tuner^26^ and the TwinsUK training set and identified the following optimized values: Intermediate layer dimensionality = 200, learning rate = 0.001, and Kullback-Leibler (KL) divergence weight = 0.01. Note that despite our hyperparameter choices, other optimal hyperparameters exist and can be chosen through Keras Tuner. Using these hyperparameters, we then optimized *d* by calculating the reconstruction MSE of the correlation matrix (CM-MSE) of metabolites for *d* = 5, 10, 15, 18, 20, 30, 40, 60, 80, 100, 120, 160, and 200 on the TwinsUK test set. Our final model consisted of a 217-dimensional input/output layer (the number of metabolites in our datasets), a 200-dimensional intermediate layer, and an 18-dimensional latent layer. For all sample encodings in the study, we used their respective μ values.

All models were computed on a deep learning-specific virtual machine running on Google Compute Engine with two NVIDIA Tesla K80 GPU dies and 10 virtual CPUs.

### 4.5. PCA embedding and reconstructions

We used PCA with *d* = 18 latent dimensions as a baseline model. On the mean-centered TwinsUK training set data matrix with *n* = 3,947 samples (rows) and *k* = 217 metabolites (columns), we calculated the rotation matrix ***Q***, a *k x k* matrix of eigenvectors ordered by decreasing magnitudes of eigenvalues. To embed a new *m* x *k* dataset ***X*** with *m* samples into the *m x d* PCA latent space ***A***, we first calculated ***XQ*** = ***A*** and subsetted to the first *d* columns, denoted by ***A***∗_*,d*_. To simulate the process of encoding and decoding in PCA for dataset ***X***, we calculated the reconstructed dataset as XJ = ***A***∗_*,d*_**Q** ^−1^_d,∗._

### 4.6. Model assessments

We assessed our PCA and VAE models using sample reconstruction mean squared error (MSE) and metabolite-wise correlation matrix MSE (CM-MSE). We calculated CM-MSE by first computing the metabolite-wise correlation matrix of an input dataset and reconstructed input dataset. Afterwards, we calculated the MSE between the upper triangular matrix of the two symmetric correlation matrices.

To calculate a confidence interval for both MSE and CM-MSE between our input and reconstructed data, we randomly sampled the same samples with replacement from the two datasets and then calculated MSE and CM-MSE. We performed this for 1,000 iterations.

### 4.7. Model interpretation

In order to interpret each latent dimension for our VAE and PCA models, we calculated Shapley Additive Global Importance (SAGE) values^30^ for metabolites, sub-pathways, and super-pathways. Briefly, SAGE is a model-agnostic method that quantifies the predictive power of each feature in a model while accounting for interactions between features. This is achieved by quantifying the decrease in model performance when combinations of model variables are removed. Since there are exponentially many combinations of variables, the current approach is to sample the feature combination space sufficiently. For each of the tested combinations, a loss function, such as MSE, is used to quantify the decrease in performance compared to the model output (here each VAE or PCA latent dimension) computed using the full model. Then, the mean of all MSEs is calculated, which represents the contribution of the model variables to a latent dimension. To calculate pathway-level SAGE values, metabolites were grouped into pathways and each pathway was treated as a single variable. For each of our VAE and PCA models, we ran SAGE using our TwinsUK test set with default parameters, e.g., marginal sampling size of 512, as suggested by Covert, et al. (2020)^30^. We used the SAGE code from https://github.com/iancovert/sage.

## Data availability

The type 2 diabetes (QMDiab), schizophrenia, and acute myeloid leukemia (AML) datasets to used in this study are available upon request from the authors and will be shared publicly when the peer-reviewed version of the manuscript is published. The TwinsUK dataset can be accessed via https://twinsuk.ac.uk/resources-for-researchers/access-our-data/uponrequest.

## Code availability

Codes used in this study are available at the GitHub repository https://github.com/krumsieklab/mtVAE

## Acknowledgements

The construction of the deep learning models was supported by Google Cloud. This work was also partially supported by the National Cancer Institute of the National Institutes of Health under the awards U10CA180820, UG1CA189859; and by the National Institute of Aging of the National Institutes of Health under award 1U19AG063744. TwinsUK is funded by the Wellcome Trust, Medical Research Council, European Union, the National Institute for Health Research (NIHR)-funded BioResource, Research Facility and Biomedical Research Centre based at Guy’s and St Thomas’ NHS Foundation Trust in partnership with King’s College London.

## Contributions

L.C., E.P., and H.F. provided the acute myeloid leukemia data. H.A. provided the schizophrenia data. K.S. provided the QMDiab data. D.P.G. and J.K. conceived of and designed the research study. D.P.G. performed model training, experiments, and analyzed all data and results. A.S. processed the datasets. D.P.G., A.S., and J.K. wrote the manuscript. All authors gave final approval to publish.

## Extended data

**Extended Data Figure 1.**
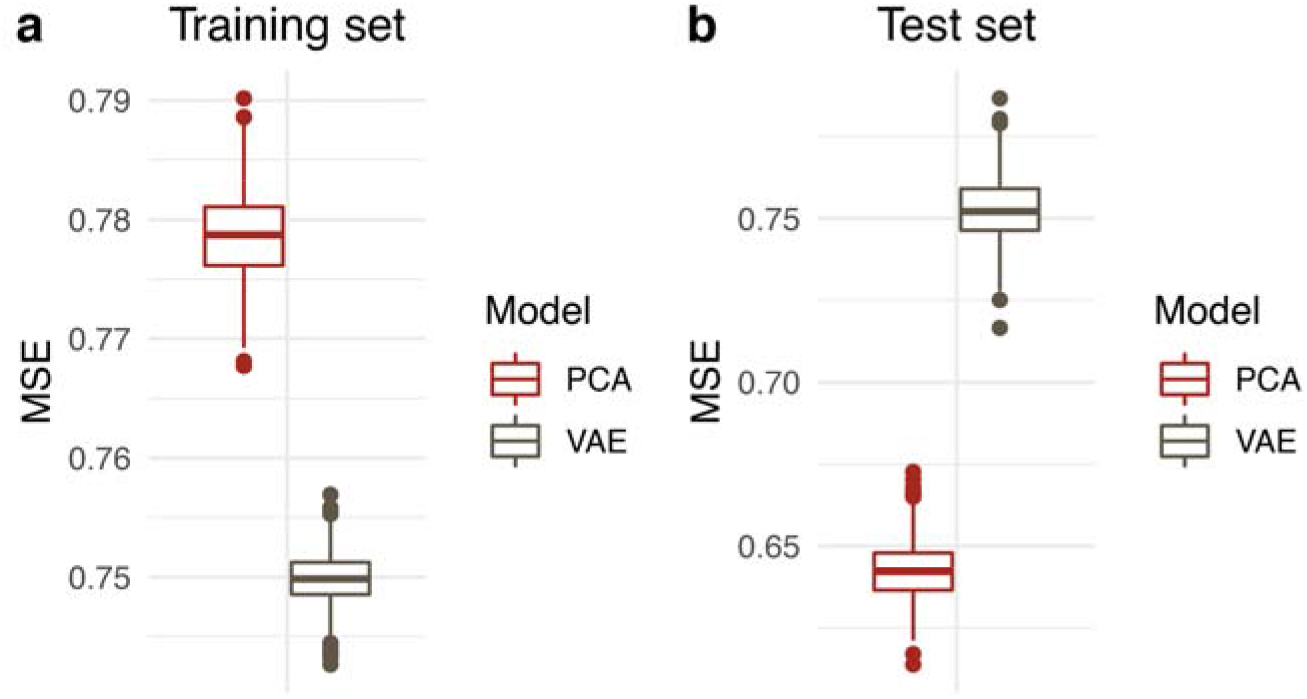
Sample reconstruction MSE. **a**, TwinsUK training and, **b**, test set sample reconstruction MSE for latent dimensionality *d* = 18. VAE has a lower reconstruction error in the training set. However, PCA has a lower reconstruction MSE in the training set, implying that PCA performs better at sample reconstruction.

**Extended Data Figure 2.**
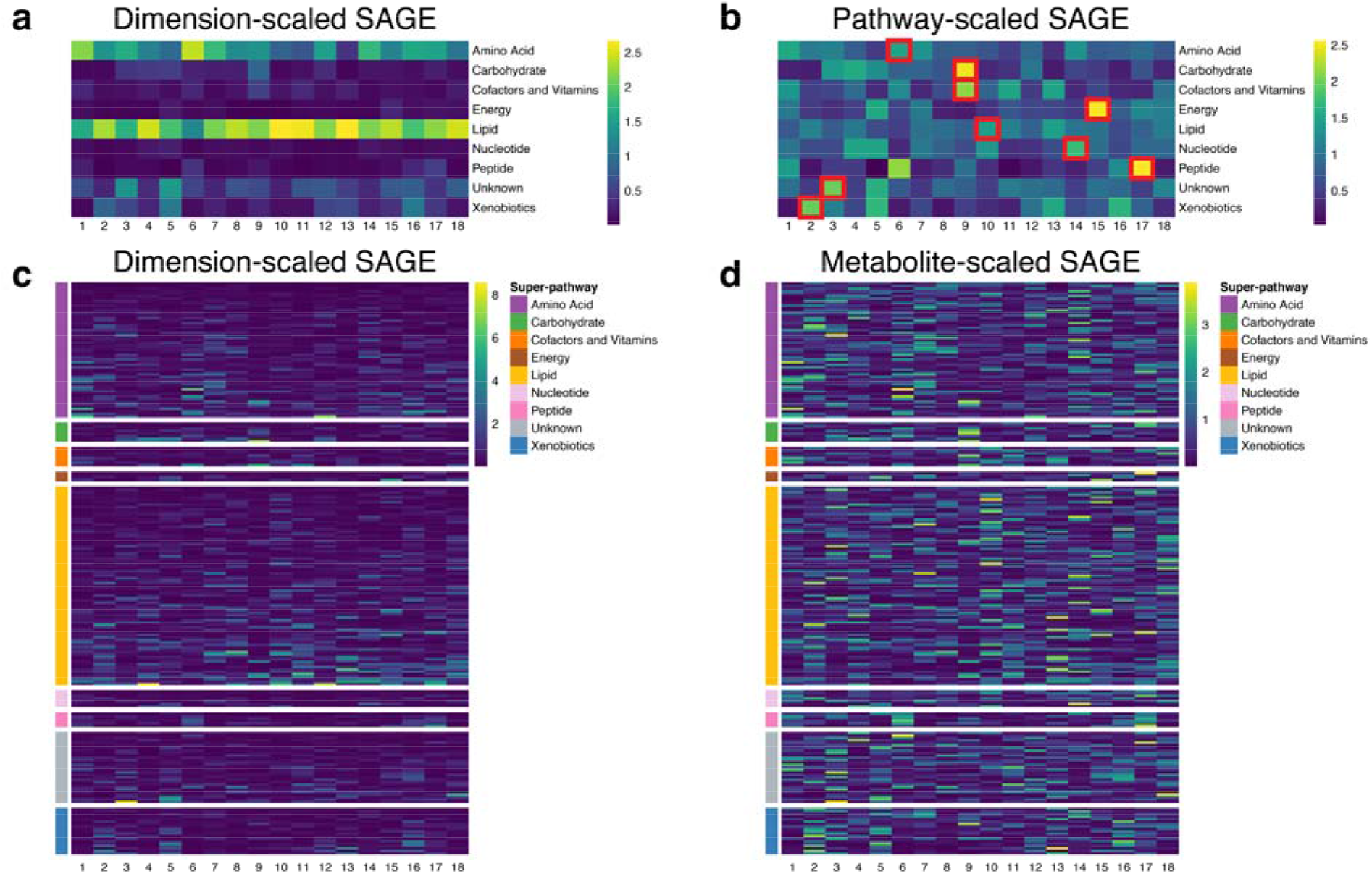
Super-pathway and metabolite-level SAGE values for the VAE latent dimensions. **a**, SAGE values were scaled by dimension, i.e. set to standard deviation 1 for each column in the matrix. This highlights pathways that contribute the most to each dimension. Lipid and amino acid super-pathways showed the highest values for most dimensions, which can likely be attributed to the high number of metabolites in those pathways. **b**, SAGE values were scaled by pathway, i.e. set to standard deviation 1 for each row in the matrix. This highlights dimensions that contribute to a pathway the most. Taking into consideration the largest scaled SAGE values per pathway (red square marks), all super-pathways are represented by unique dimensions. **c**, Absolute metabolite SAGE were scaled by dimension, i.e. set to standard deviation 1 for each column in the matrix. This highlights metabolites that contribute the most to each dimension. Metabolites in the lipid and amino acid super-pathways showed the highest values for the majority of the dimensions. **d**, Absolute metabolite SAGE values were scaled by metabolite, i.e. set to standard deviation 1 for each row in the matrix. This highlights dimensions that contribute to a metabolite the most. Each dimension has a specific metabolic signature. The combination of these metabolites of a dimension outlines the distinct cellular mechanisms a dimension encodes.

**Extended Data Figure 3.**
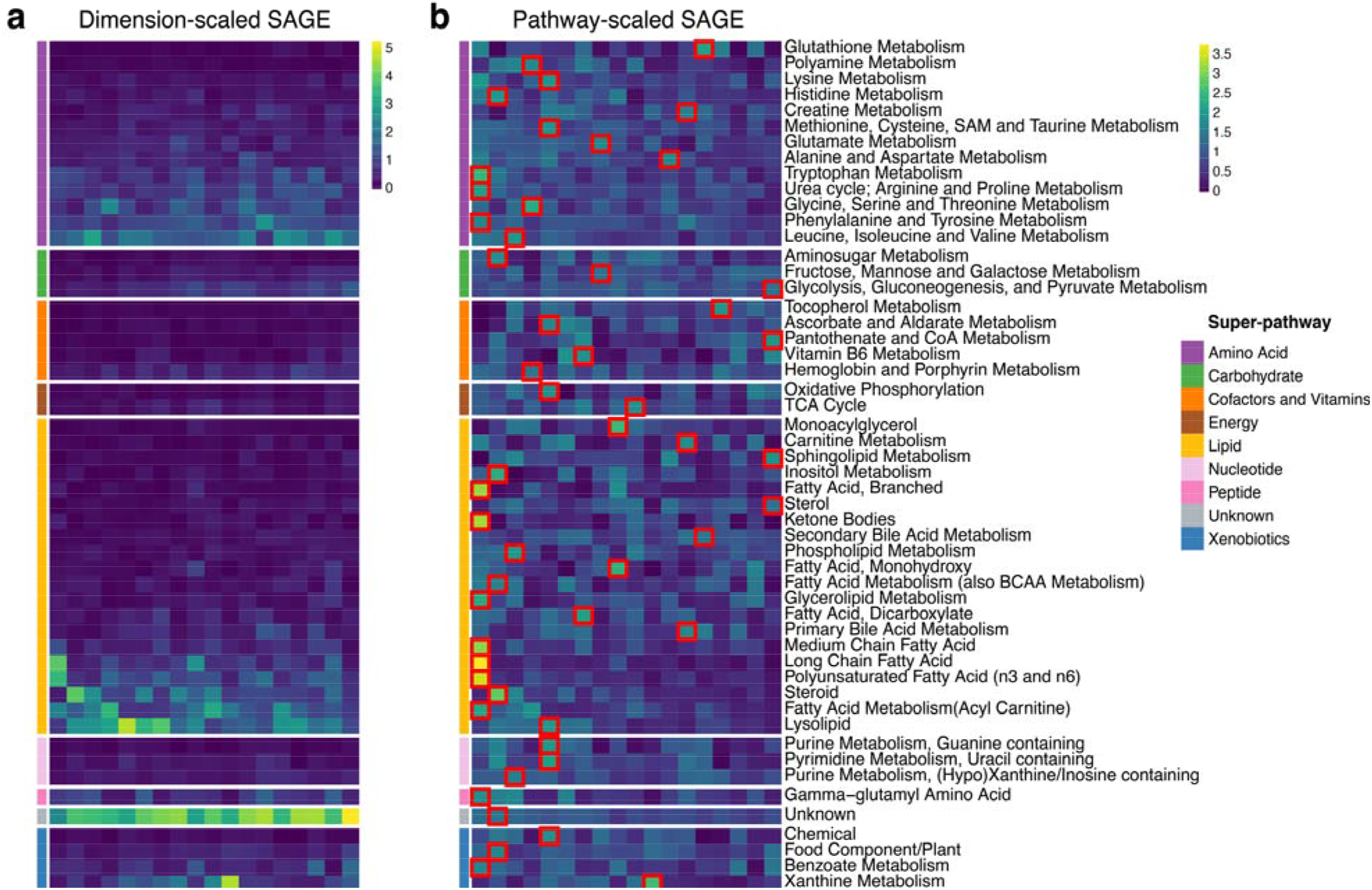
Sub-pathway-level SAGE values for the PCA latent dimensions. **a**, SAGE values were scaled by dimension, i.e. set to standard deviation 1 for each column in the matrix. This highlights pathways that contribute the most to each dimension. Lipid, unknown, and amino acid super-pathways showed the highest values for most dimensions, which can likely be attributed to the high number of metabolites in those pathways. **b**, SAGE values were scaled by pathway, i.e. set to standard deviation 1 for each row in the matrix. This highlights dimensions that contribute to a pathway the most. Taking into consideration the largest scaled SAGE values per pathway (red square marks), sub-pathways concentrate on the first 3 dimensions, especially on dimension 1. Other dimensions have primarily unrelated sub-pathways.

**Extended Data Figure 4.**
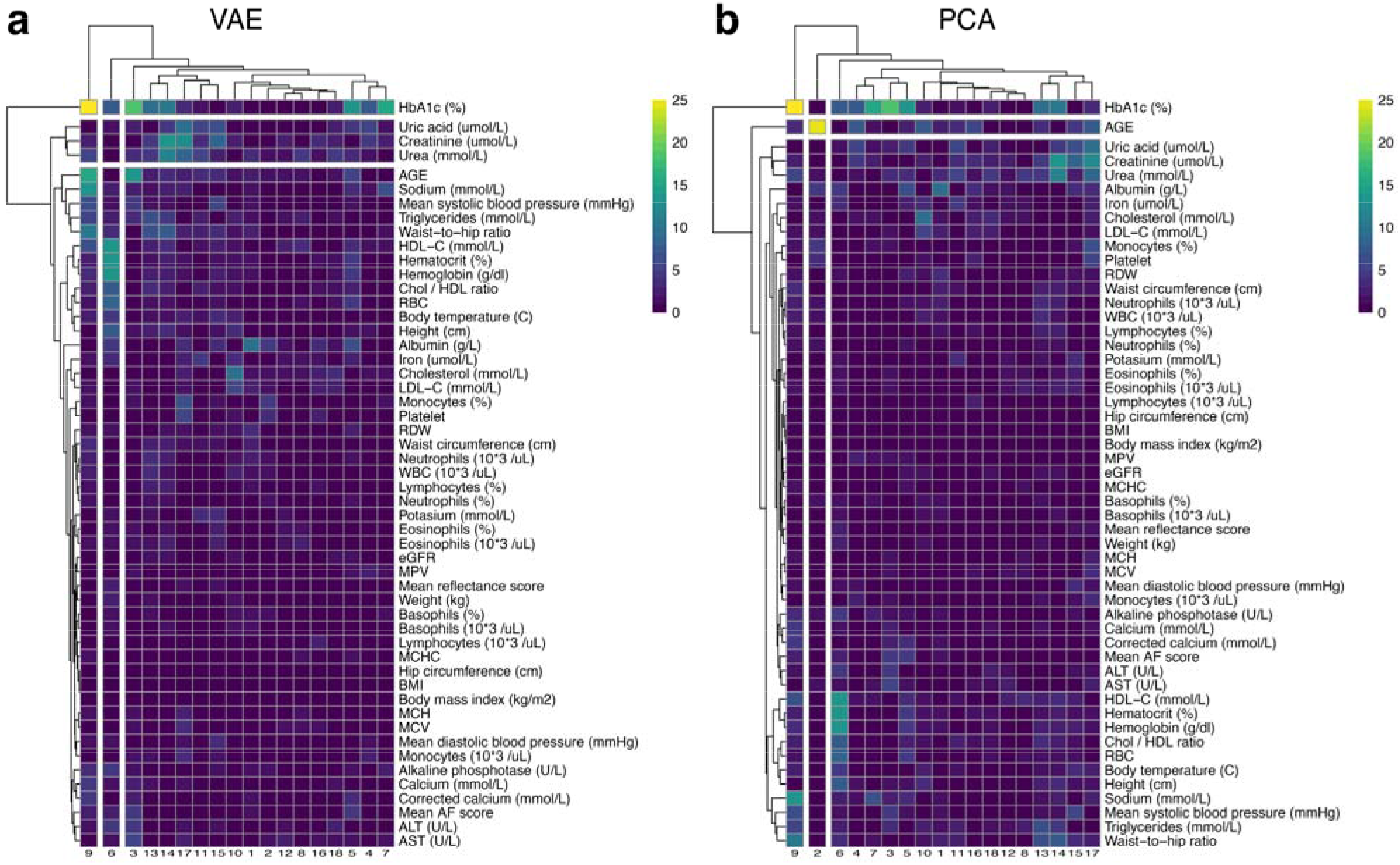
Type 2 Diabetes clinical variable associations with VAE and PCA latent dimensions. Association heatmap between for **a**, VAE and **b**, PCA latent dimension values. Each latent dimension is associated with different combinations of clinical variables. Both VAE dimension 9 and PCA dimension 16, which associate with QMDiab diabetes groups, strongly associate with HbA1c (%). VAE dimension 9 HbA1c association p = 5.6×10^−56^, PCA dimension 16 HbA1c association p = 1.1×10^−30^.

**Extended Data Figure 5.**
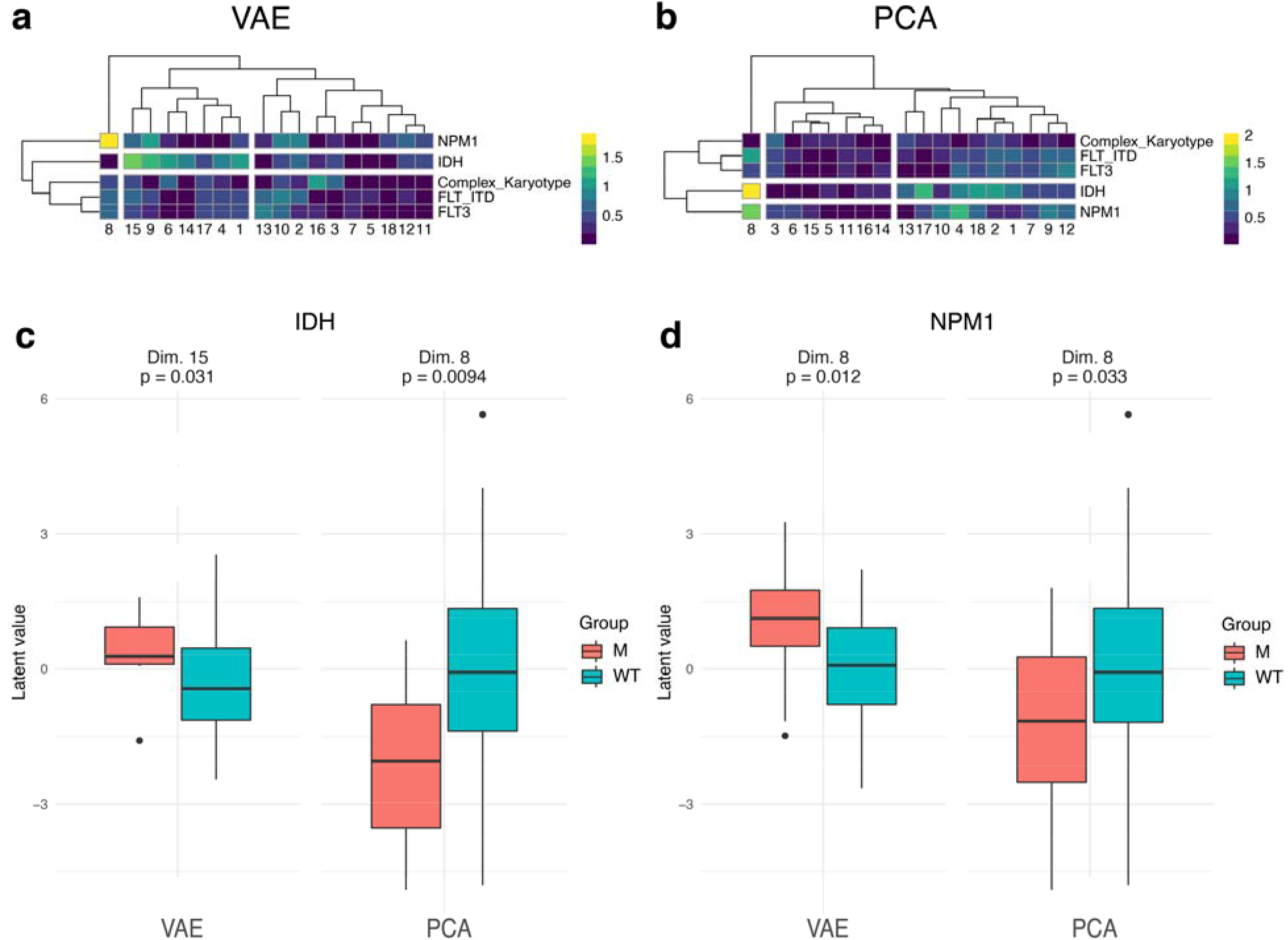
AML mutation profile and latent dimension associations. Our dataset initially contained 21 AML-related mutations: AML1-ETO, ASXL1, CBF, CEBPa, DNMT3A, EVI1, FLT-ITD, FLT3, IDH1, IDH2, KIT, KRAS, MLL, NPM1, NRAS, PHF6, PTEN, RUNX1, TET2, TP53, WT1. To ensure adequate statistical power, we selected mutations with at least 10 samples per group, i.e. mutant or wildtype. This criterion retained 4 mutations and “complex karyotype” for our final statistical analysis. **a**, VAE latent dimensions association heatmap. **b**, PCA latent dimension association heatmap. IDH and NPM1 show the strongest associations to the latent dimensions. **c**, boxplot of VAE and PCA latent values for IDH. PCA dimension 8 associates stronger with IDH. **d**, boxplot of VAE and PCA latent values for NPM1. VAE dimension 8 associates more with NPM1. Color bars for **a** and **b** are -log_10_(p-value). For **c** and **d** M = mutant, WT = wildtype.

## Extended Data information

### Extended Data Tables

Extended Data Tables 1-3

